# Basal forebrain rhythmicity is modulated by the exploration phase of novel environments

**DOI:** 10.1101/2020.01.11.902742

**Authors:** Diogo Santos-Pata, Paul FMJ Verschure

**Affiliations:** SPECS, Institute for Bioengineering of Catalonia, Barcelona, 08028, Spain; Institució Catalana de Recerca i Estudis Avançats (ICREA), Barcelona, 08010, Spain

**Author notes:** **Correspondence** Paul FMJ Verschure, SPECS, Institute for Bioengineering of Catalonia, Barcelona, 08028, Spain. SPECS, Institute for Bioengineering of Catalonia, Barcelona, 08930, Spain (Paul FMJ Verschure).

**Keywords:** Basal forebrain, *Learning*, Hippocampus, Theta-Gamma

## Abstract

Acquaintance to novel environments requires the encoding of spatial memories and the processing of unfamiliar sensory information in the hippocampus. Cholinergic signaling promotes the stabilization of hippocampal long-term potentiation (LTP) and contributes to theta-gamma oscillations balance, which is known to be crucial for learning and memory. However, the oscillatory mechanisms by which cholinergic signals are conveyed to the hippocampus are still poorly defined. We analyzed local field potentials from the basal forebrain (BF), a major source of cholinergic projections to the hippocampus, while rats explored a novel environment, and compared the modulation of BF theta (4-10Hz) and gamma (40-80Hz) frequency bands at distinct stages of spatial exploration. We found that BF theta and gamma display learning stage-related rhythmicity and that theta-gamma coupling is stronger at the later stages of exploration, a phenomenon previously observed in the hippocampus. Overall, our results suggest that the BF-hippocampal cholinergic signaling is conveyed via the stereotypical oscillatory patterns found during mnemonic processes, which questions the origins of the learning-related rhythmic activity found in the hippocampus.

**KEY-POINTS:**

- Basal forebrain theta oscillations decrease their strength in function of exploration time, as observed in the hippocampus.
- BF gamma ripples (bursting events) are longer after learning.
- BF Theta-gamma coupling increases after initial spatial exploration, suggesting BF cross-frequency coupling relation to the learning stage.

## 1 INTRODUCTION

The formation of spatial representations is generally attributed to the hippocampal circuitry where a myriad of cell types encoding distinct aspects of one’s surroundings allow to decode current position and future trajectories (Nadel, 1991). The hippocampal cognitive-map is thought to be formed via a combination of environmental/sensory and self-generated analog signals, encoded both in single-unity activity (Moser et al., 2008) and population-level oscillations (O’Keefe, 1993), with the end result of serving ongoing navigation and spatial decision-making (Buzsáki and Moser, 2013; Wikenheiser and Redish, 2015). Hippocampal theta and gamma oscillatory activity and synchrony play a role in learning and memory (Düzel et al., 2010; Tort et al., 2009), and hippocampal theta frequency has been shown to decrease logarithmically in function of training time, relating theta oscillatory components with the early learning stages (Pan and McNaughton, 1997). Coincidentally, selective silencing of septal cholinergic neurons (within the basal forebrain) results in hippocampal theta rhythmicity decreases, which might explain the cognitive anomalies associated with dementia (Sainsbury and Bland, 1981; Hasselmo, 2000). Indeed, Alzheimer’s disease (AD) patients show neuropathologies such as degeneration of cholinergic BF neurons, suggesting a relationship between BF cholinergic hypofunction and cognitive impairments (Auld et al., 2002; Ballinger et al., 2016).

Septal nuclei projections to the hippocampus are known to be a source component of hippocampal theta waves (Stewart and Fox, 1990). The rhythmic firing of cholinergic and GABAergic septal-hippocampal projections, together with intra-hippocampal excitation, lead to pronounced hippocampal theta waves (Stewart and Fox, 1990; Colom, 2006). Furthermore, learned spatial information encoded in the hippocampus was observed to be transmitted to the lateral septum via theta-dependent neuronal sequences, emphasizing the role of theta waves in mediating hippocampus-BF communication (Tingley and Buzsáki, 2018). Together with the observations that strong hippocampal theta rhythm occurs during the early learning stages (Düzel et al., 2010; Sakimoto and Sakata, 2014), it is also known that hippocampal theta elicits neuronal excitatory postsynaptic potentials (EPSPs) so that it boosts synaptic plasticity (Huerta and Lisman, 1993). In addition, cholinergic projections originating in the BF modulate hippocampal theta and gamma oscillations, and the excitation-to-frequency transduction (discharge) of basal forebrain cholinergic neurons burst when oscillatory theta amplitude is maximal (Lee et al., 2005), suggesting that cholinergic signaling is involved in learning and memory (Dannenberg et al., 2015; Huerta and Lisman, 1993).

Decreases in rodent hippocampal theta power during learning have been observed in memory retention tasks (Sakimoto and Sakata, 2014), and human hippocampal fMRI voxel activity decreases at later stages of spatial learning (Brodt et al., 2016). Similarly, BF beta-frequency oscillations (15-35Hz) strength is modulated over time while rats perform an associative-reinforcement task (Quinn et al., 2010), with beta peaks displaying a greater amplitude at early learning stages.

Thus, in light of the relationship between cholinergic signaling, originated within the BF nuclei, and the learning stages mediating hippocampal rhythms, it is still unclear what are the oscillatory properties of BF during spatial exploration. Here, we hypothesize that the hippocampal theta oscillatory strength observed during learning stages is accompanied by equally increased theta strength in its cholinergic origins (i.e. Basal forebrain). If that was the case, learning-related theta waves observed in the hippocampus would indicate a broader communication channel used to engage the cholinergic pathway. In order to assess if and how BF oscillations are affected by learning, we analyzed a dataset (Nair et al., 2018a,b) of BF LFPs from rats (n=6) performing spatial navigation within a novel arena at distinct stages of exploration (with navigational sessions lasting 673+/−48sec, mean/std).

## 2 RESULTS

Our primary question focused on the oscillatory properties of BF during spatial learning. We split the dataset into *early* and *late* exploration phases by extracting both LFP and behavioral signals from the first and third quartile of the exploration duration (172+/−10 sec, mean/std, Fig1-A). As novelty detection and early exploration are thought to be expressed behaviorally (Lisman and Otmakhova, 2001; Jeewajee et al., 2008; Vago and Kesner, 2008), we analyzed the movement signal profiles of each animal during early and late phases. Four out of 6 animals displayed higher movement (displacement in the 3-axis) at early phases (t-test, p<0.05). At the population level, however, we did not observe a significant difference in the movement signal for early (317 +/−169, mean+/−std) compared with late (278 +/−150, mean+/−std) exploration phases (Fig1-B).

**FIGURE 1.**
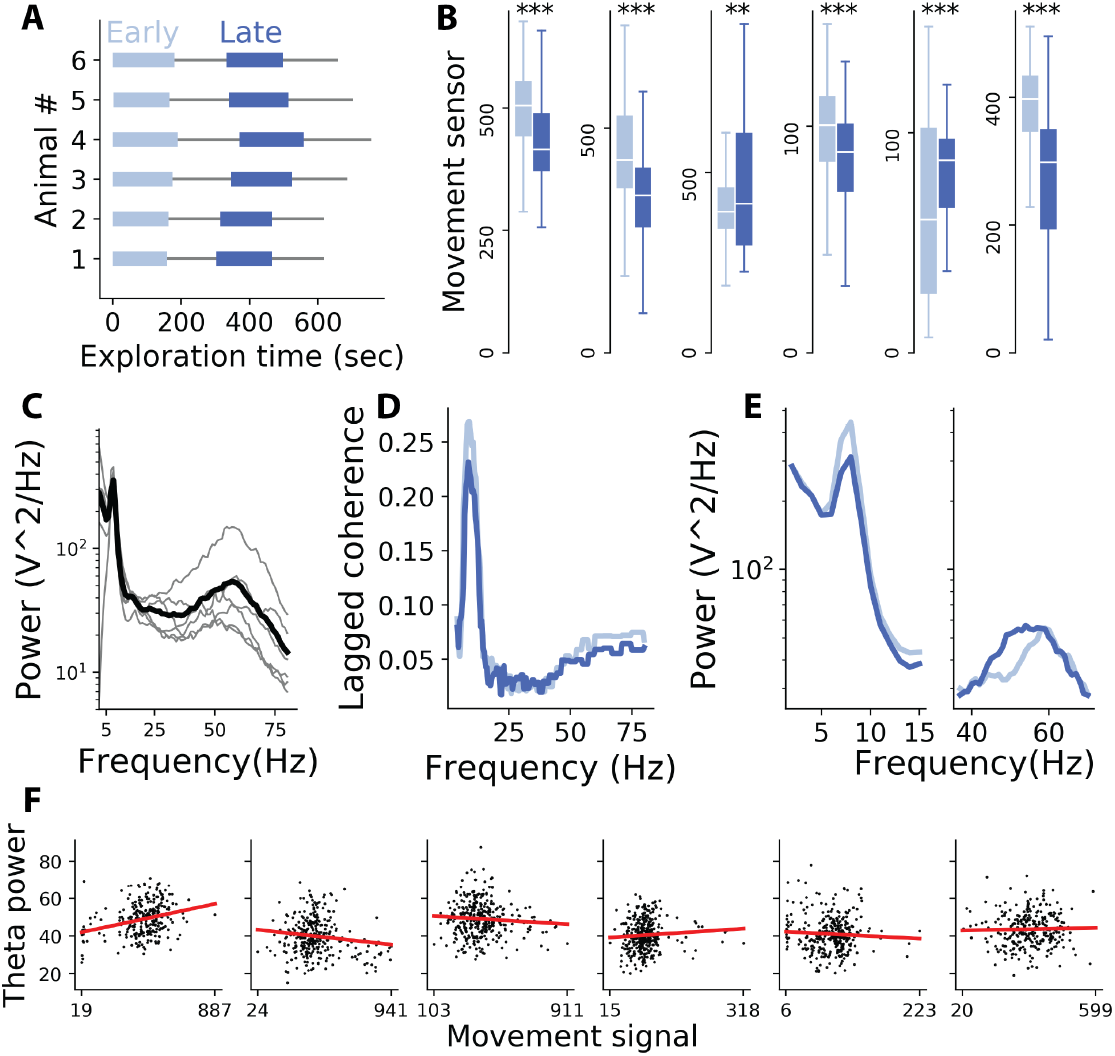
Behavioral and LFP characterization of early and late spatial exploration. **A** Exploration times for each individual (grey horizontal lines, 673+/−48 sec, mean/std), and the periods of ‘Early’ and ‘Late’ exploration (light and dark blue, respectively, 172+/−10 sec, mean/std). **B** Distribution of movement sensor values during early and late exploration periods (t-test, **<0.01, ***<0.001). **C** Power spectral density (Welch) extracted from BF LFPs recorded during the entire exploration session for each animal (grey lines), and the average PSD (black line). **D** Strong theta rhythmicity (lagged coherence) for early (peak=9.0Hz) and late (peak=8.5Hz) exploration phases (4-80Hz frequency interval spaced at 0.5Hz). **E** BF spectral power (as in B) for theta (left) and gamma (right) for early and late exploration phases. Note the increased theta power at early exploration and the increased gamma power at late exploration phases. **F** Theta power (instantaneous amplitude) averaged (mean) for each 2-second time window in function of the animal movement revealed no significant correlation.

Afterward, we computed the basal forebrain oscillatory strength of individual frequencies along with the alpha to gamma range (1-80Hz) and observed the stereotypical theta and gamma frequency bands pronounced density characteristic of hippocampal LFPs (Fig1-C). A result further confirmed by the strong theta frequency rhythmicity resulting from lagged coherence analysis (Fig1-D). Moreover, the splitting of BF LFPs in early and late exploration suggested that theta and gamma frequency bands display a distinct spectral density modulation accordingly with the learning phase, that is, stronger theta at the early stage and stronger gamma at later stages (Fig1-D and E).

Because hippocampal theta oscillatory components are known to be modulated by the locomotory speed in both rats (McFarland et al., 1975; Santos-Pata et al., 2017) and humans (Ekstrom et al., 2005), the higher theta (and lower gamma) amplitude observed in the early exploration phases could be a cofound of higher movement signals observed also in early exploration (Fig1-B, D and E). As a control for the possible movement-related evoked theta, we analyzed BF theta oscillatory amplitude in relation to the animal movement signal. None of the six animals revealed a movement-theta trend surviving the correlation confidence interval threshold (ci=0.1, Pearson-r test, Fig1-F), therefore discarding the effects of locomotion in modulating BF theta oscillations.

In order to assess how the oscillatory strength of BF theta waves is modulated by the learning stage, we extracted the instantaneous amplitude of the Hilbert-transformation in the 4-10Hz range for each 2-second time window belonging to early and late exploration phases (Fig2-A). Individually, all six animals presented a stronger theta amplitude at the early learning phase (t-test, p<0.05), which was confirmed by a population-level analysis where theta amplitude at early (43.97+/−4.06uV, mean+/−std) and late (38.65+/−3.03uV, mean+/−std) stages were significantly different (t-test, statistic=2.35, p=0.04, Fig2-B). The learning phase-dependent theta modulation was further computed using the Hilbert transformation (filter length of 3 cycles of 4Hz cutoff frequency) and averaged across the individual theta cycles analytic amplitude (Fig2-C). Again, the averaged cycle analytic theta was higher at early (47.09+/−4.01uV, mean+/−std) compared to late (41.89+/−3.87uV, mean+/−std) exploration phases and showed a significant decrease in the oscillatory strength (t-test, statistic=2.26, p=0.04), a result that could not be explained by the relation between the animal locomotory behavior and theta amplitude (Fig1-F).

**FIGURE 2.**
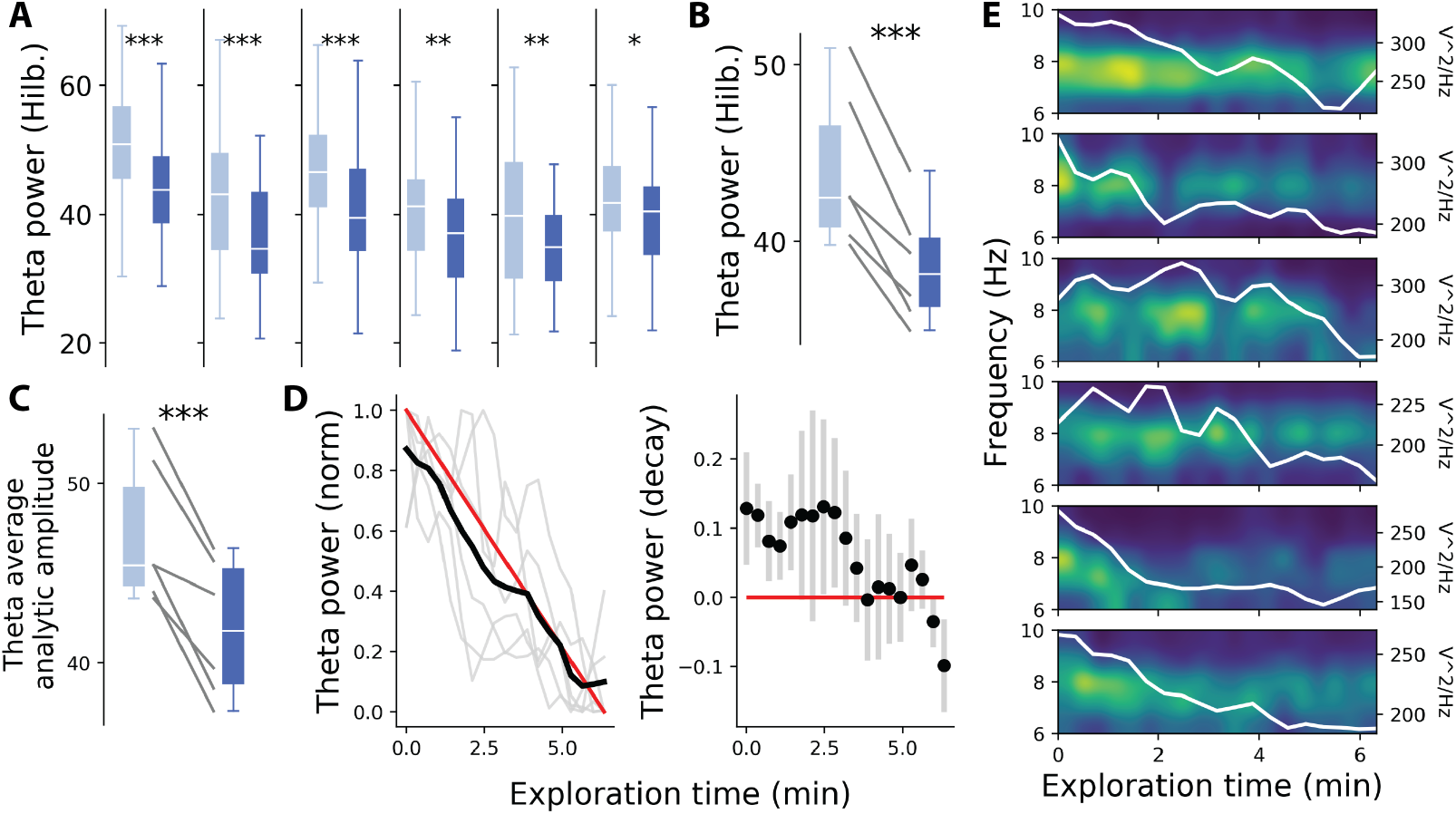
Theta (4-10Hz) and gamma (40-80Hz) BF spectrum are modulated by the exploration phase and animal movement. **A** Theta amplitude is higher during the early exploration phase (p < 0.05*, 0.01**, 0.001***, T-test). **B** Population-level comparison of early (43.97+/−4.05, mean/std) vs late (38.65+/−3.03, mean/std) theta amplitude (T-test related samples, statistic=7.02, p<0.001, effect size = 1.48). **C** Theta power extracted from each cycle analytic amplitude at early (47.09+/−4.01, mean/std) and late (41.89+/−3.87, mean/std) exploration confirmed the power decreases observed in A and B (T-test related samples, statistic=6.08, p=0.001, effect size=1.31). **D** Left: Individual (grey lines) and averaged (black line) normalized theta amplitude along the exploration session, superimposed by a fictional linear decay (red line). Right: Difference between the linear decay function and the normalized theta power. **E** LFP spectrograms for each rat during navigation (6.2min), clipped to 6-10Hz theta band. Decreases in theta power density (20sec bin averaged) along the exploration session (white line).

Having assessed the changes in theta strength from early to late exploration phases, we next asked how are such amplitude decreases reflected along the entire exploration time (673+/−48 sec, mean/std). To assess the decay profile of theta strength throughout exploration, we re-computed the power spectral density (Welch’s method) of BF LFPs in the theta range (4-10Hz) and averaged it in 20 seconds time bins along with the first three quartiles of the exploration session (Fig2-D and E). Visual inspection of the oscillatory spectrogram in the high theta range (6-10Hz) for each animal depicted a linear trend in the decay function of oscillatory strength over time (Fig2-E). Indeed, the normalized theta amplitude was significantly anti-correlated with exploration time (Pearson-R test, r=−0.99, p<0.001, confidence interval: min=−0.996, max=−0.975; Polynomial coefficients degree-one fit = −0.1307× + 0.8562, Fig2-D left). We observed, however, a stairs-shaped trend along the exploration time when computing the difference between a linear decrease function and the normalized theta amplitude for each animal, suggesting a resistance against zero strength pronounced after 2.5min of exploration (Fig2-D right), which might be informative of the temporal dynamics of learning during early exploration.

It has been previously seen that hippocampal gamma power increases with memory load (van Vugt et al., 2010) and that those same increases contribute to memory consolidation immediately after learning (Pu et al., 2018). Thus, if BF theta waves reflect aspects of cognition, higher frequencies of the same local field should also participate in spatial learning. Similarly to those hippocampal observations, we also found a significant increase from BF gamma amplitude at early (33.05+/−10.8, mean/std) to late (35.95+/−10.50, mean/std) exploration phases (t-test, statistic=−4.38, p=0.007, effect size = −0.27), suggesting that BF gamma waves are also modulated by the learning stage (Fig3-B).

**FIGURE 3.**
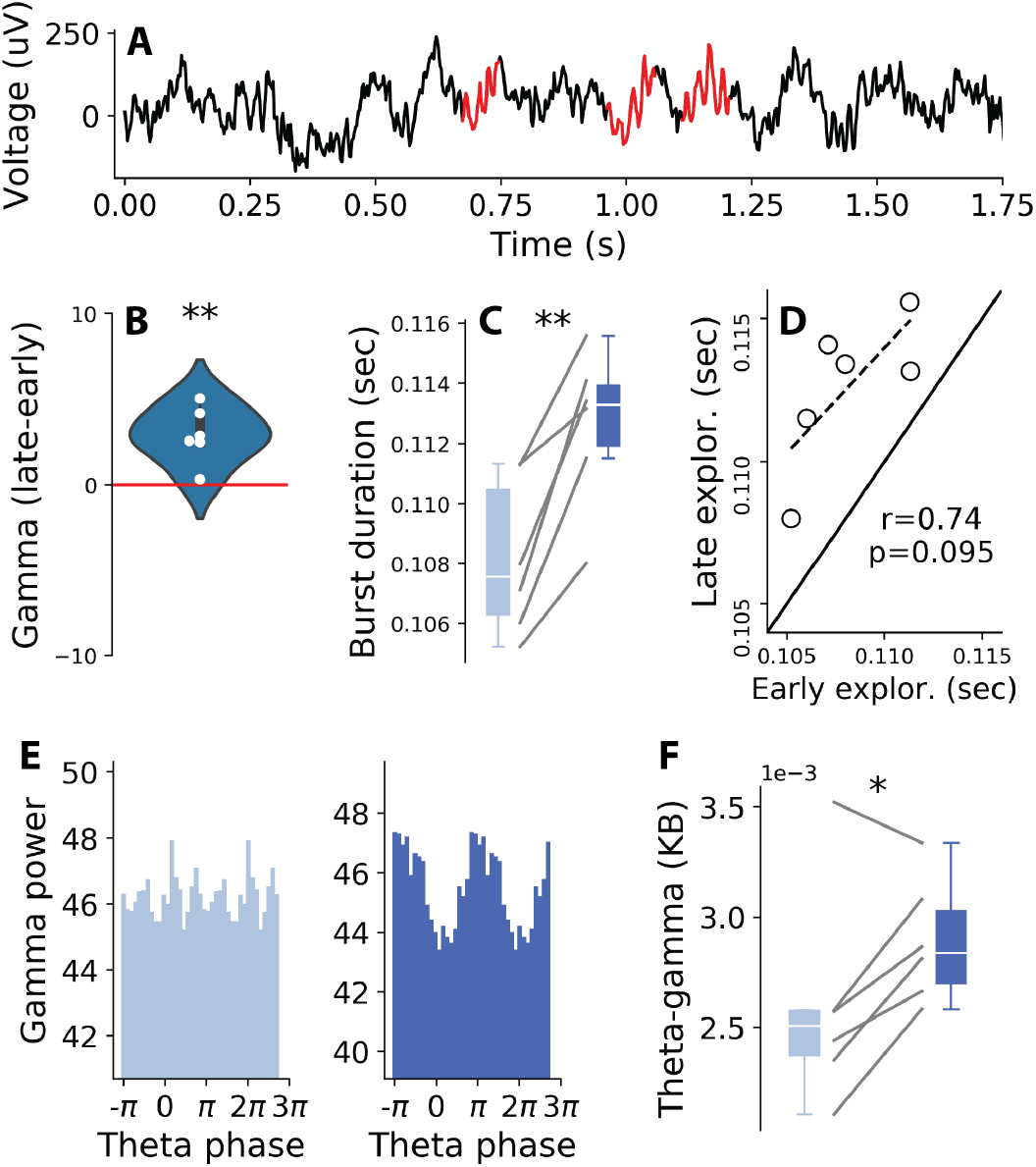
Exploration phase modulates gamma activity and theta-gamma coupling **A** An example of raw LFP trace (black line) with highlighted segments of gamma (40-80Hz) bursting activity (red lines), computed using the dual amplitude threshold algorithm (see methods section). **B** BF gamma amplitude is higher (35.95+/−10.82 mean+/−std) at later compared to the early (33.05+/−10.50 mean+/−std) stages of exploration (related sample T-test; statistic=−4.38, p=0.007, effect size=−0.27). **C** Gamma bursting activity is significantly shorter at early (0.108+/−0.002 mean+/−std) than late (0.112+/−0.002 mean+/−std) exploration phases (t-test: statistic=−1.86, p=0.002). **D** Individual bursting duration for early and late exploration phase (Pearson test; r=0.74, p=0.095). **E** Example of the averaged gamma power distribution along with theta phase (one rat, id=2), at early (left) and late (right) exploration phases. **F** Cross-frequency coupling computed by the divergence (Kullback-Leibler) score for early (2.5e-3+/−0.4e-3 mean+/−std) and late (2.8e-3+/−0.2e-3 mean+/−std) learning stages (related sample T-test; statistic=−2.77, p=0.03, effect size=−0.81).

The role of BF oscillatory bursting in the beta-band (15-30Hz) has been linked with associative memory (Quinn et al., 2010) and the duration of hippocampal gamma ripples has been recently observed to prolongate in tasks involving spatial memory (Fernández-Ruiz et al., 2019). To answer whether BF gamma bursting duration is also related to the distinct learning stages, we quantified BF gamma bursting duration at early and late exploration phases by applying a dual amplitude threshold algorithm to BF LFPs (Cole et al., 2019). Burst identification was set with a cutoff threshold interval of 1-1.5std of the normalized (z-scored) LFP signal filtered in the low gamma band (40-60Hz) and event duration was extracted by the time interval between the upward crossing of 1.5std until the downwards crossing of 1std (Fig3-A). Interestingly, gamma burst events duration at an early stage (0.108+/−0.002sec, mean/std) were significantly shorter than the ones found during the late stage (0.113+/−0.002sec, mean/std) (t-test; statistic=−5.75; p=0.002, effect size =−1.86, Fig3-C), a small but sufficiently longer burst prolongation to include an extra high-frequency oscillatory cycle, characteristic of hippocampal sharp-wave ripples (Fernández-Ruiz et al., 2019). Notably, the exploration period did not greatly affect each animal BF bursting, but rather slightly modulated burst event duration (Fig3-D).

Having found changes in the rhythmicity of both theta and gamma waves, we next focused on their interaction. Theories of hippocampal cross-frequency coupling highlight the role of theta waves in carrying item related information expressed in the form of gamma cycles, which has been related to working memory and cognitive load (Lisman and Jensen, 2013). In light of such a coding scheme, together with the findings that hippocampal gamma amplitude is more strongly modulated by the theta oscillatory phase at later stages of learning (Tort et al., 2009), we asked whether also BF theta-gamma coupling is affected by the exploration stage. We extracted the instantaneous theta-phase and gamma-amplitude of each 2-second time window belonging to early and late exploration phases and calculated the cross-frequency coupling against a uniform distribution using the Kullback-Leibler divergence score (see methods). Concurrently with the hippocampal observations, we also found a significant increase in the theta-gamma modulatory index from early (0.0025+/−0.0004, mean/std) to late (0.0029+/−0.0002, mean/std) exploration phases (t-test; statistic=− 2.77; p=0.03, effect size =−0.82), which supports the idea that BF cross-frequency coupling also plays a role in mnemonic processes (Fig3-E and F).

## 3 DISCUSSION

Great emphasis has been put on the functional role of the hippocampus in spatial learning, memory and novelty detection. At the physiological level, hippocampal theta and gamma oscillations have been analyzed in the context of sensory encoding, associative learning and spatio-temporal representations at multiple scales (e.g. phase precession (O’Keefe and Recce, 1993)). The responses of different frequency bands along the learning stages has led to the general agreement that the hippocampus is actively involved in setting the oscillatory components needed to facilitate learning. However, at the neuromodulatory level, it is known that basal forebrain cholinergic projections to the hippocampus facilitate neural plasticity (Dannenberg et al., 2015; Huerta and Lisman, 1993; Lee et al., 2005) and that the cognitive deficits manifested in AD (often associated with hippocampal damage) originate in the BF due to cholinergic hypofunction (Auld et al., 2002). Thus, we questioned whether the oscillatory specifics of learning originate in the hippocampus or, instead, are already present in presynaptic sites of the cholinergic system, such as BF populations. To answer this question, we analyzed local field potentials of rats during novel-environment exploration and assessed BF oscillatory properties at distinc exploration phases. Analogous with the spectral density seen in the rodent hippocampus during active navigation, we also observed a strong oscillatory amplitude in theta (4-10Hz) and gamma (40-80Hz) frequency bands in the basal forebrain of rats exploring a novel environment (Fig1-C-E). Interestingly, we did not find a relation between the locomotory speed and theta strength as found in the hippocampus, suggesting that BF oscillations are agnostic to overt locomotory behavior (Fig1-F).

With respect to learning, our results indicate that both BF theta and gamma waves manifest their amplitude profiles distinctly in early and late exploration phases (Fig1A-C and Fig2-C and D), proposing that BF actively participates in the spatial learning circuitry. Moreover, we quantified the decay rate of theta and observed a linear, rather than abrupt, decrease in the oscillatory amplitude as the animal explores a novel environment, which argues in favor of continual assessment of the animal’s learning (Fig2-D and E). A piece of accompanying evidence supporting the notion that BF oscillations carry spatially-related information was the observation that gamma oscillatory bursts tend to have a longer duration at later stages of learning (Fig 3-C), a phenomenon previously quantified in the hippocampus of rats during spatial memory tasks (Fernández-Ruiz et al., 2019; Santos-Pata et al., 2019). Furthermore, we evaluated whether phase-amplitude cross-frequency (theta-gamma) coupling had a functional relevance for learning as it is expected from hippocampal oscillations (Fig3-E) (Tort et al., 2009). Unexpectedly, we observed that the amplitude of gamma was strongly carried (modulated) by theta phase at later stages of exploration, suggesting that BF theta-gamma code plays a role in spatial learning (Fig3-F). Altogether, our results highlight the engagement of BF in mediating learning and strengthen the hypothesis that cholinergic signaling to the hippocampus is conveyed via learning stage-related oscillatory patterns. The oscillatory profile of BF waves during spatial learning raises the hypothesis that BF integrates the hippocampal-cortical circuitry involved in spatial representations and contributing to overt behavior both in rodent and human navigation (Redish, 2016; Santos-Pata and Verschure, 2018). A remaining question, however, is that of how do BF and hippocampal theta oscillations interact with each other during learning. One option would be that learning-related theta strength in both areas is modulated via a common driver, and they are an echo of the biophysical interactions during learning. Another possibility is that hippocampal and BF theta waves have a strong lagged-coherence in their oscillatory phase, with the phase of BF preceding the one of the hippocampus. Such synchronization would be a solution to increase the EPSPs slope of CA1 principal cells right before the income of sensory signals expressed through gamma peaks embedded within theta cycles (Lisman and Jensen, 2013).

## 4 METHODS

### Task and participants

We analyzed BF local field potentials (LFPs) of rats (n=6) during spatial exploration of a novel environment (673+/−48 sec, mean/std) (Nair et al., 2018a,b). In order to account for learning phases, displacement (movement) and BF LFPs were grouped in earlier (first quartile) or later (third quartile) stages of exploration (172+/−10 sec, mean/std) (Fig1-B). The animal movement was extracted from a movement sensor signal, averaged for every 2-second window (see (Nair et al., 2018a) methods section). All analyses were performed using common Python packages for signal processing and statistical analyses.

### BF Local field potentials and frequency bands

Basal forebrain LFPs were obtained from chronically implanted tungsten electrodes using a wireless data logger (400Hz sampling rate) (see (Nair et al., 2018a)). Power spectral density (PSD) was computed using the ‘Welch’ method. Each animal PSD was computed for each 2-second time window and then averaged (Fig1-A). Unless otherwise stated, all analyses were performed using the NEURODSP python library (Cole et al., 2019). Analysis of theta strength decay along exploration time took into account the first 10 minutes of exploration (so that every time-bin from all animals could be included, Fig2-D). Theta and gamma oscillatory bands were chosen in the 4-10Hz and 40-80Hz range, respectively (unless otherwise specified). Theta strength modulation with respect to a linear decay function was assessed by averaging the theta band amplitude resulted from the PSD analysis for each segment along the 10 minutes (averaged and resampled in 28 bins)(Fig2-C). Next, each animal theta strength was normalized (min=0, max=1) and subtracted from a representative linear decay function (28 bins). Lagged coherence was computed using the lagged coherence function from Neurodsp (number of *cycles* = 3) and served us to confirm the strong oscillatory theta component observed with the PSD analysis (Fransen et al., 2015). Theta average analytic amplitude was extracted from each theta cycle, using the Bycycle python package (Cole and Voytek, 2019). Cycle features thresholding was set for the main oscillatory components (amplitude consistency = 0.6; periodic consistency=0.75, monotonicity= 0.8).

### Bursting activity

Oscillatory bursting events were detected using the dual-threshold burst detection function included in the Neurodsp python library (Cole et al., 2019). In short, each 2-second segment from LFPs belonging to either the early or late exploration phase was (bandpass) filtered in the low gamma band (40-60Hz) and z-scored normalized. A cutoff interval was set with a minimum (1std) and a maximum (1.5std) threshold. Bursting periods were identified by marking the moments in which the normalized filtered signal raised above the maximum cutoff threshold and maintained a sustained activity above the minimum threshold for a minimum of 3 cycles of the filtered signal (Fig3-A).

### Cross-frequency coupling

Phase-amplitude coupling was computed for each 2-second segment of LFPs in both exploration stages by extracting the instantaneous phase of theta (4-10Hz) and the instantaneous amplitude of gamma (40-80Hz). Next, we computed the statistic (mean) distribution of gamma amplitude along with the theta phase (bins=20). The modulatory index reflecting the theta (carrier) - gamma (carried) modulation was then computed using the Kullback-Leibler score as presented in (Tort et al., 2009). The averaged modulatory indexes per each animal at early and late exploration were then used for statistical analysis (Fig3-E and F).

## Abbreviations

BF: Basal forebrain
LFP: Local field potential
AD: Alzheimer’s disease

## ACKNOWLEDGEMENTS

We are grateful to Riccardo Zucca for his feedback on the early versions of the manuscript. We appreciate the efforts by the authors of the original study who made their dataset publicly available(Nair et al., 2018a,b). Without their contribution to open science this manuscript would not have been possible.

